# Histological and Metagenomic Analysis of Microbial Communities in Archaeological Human Bones

**DOI:** 10.64898/2025.12.21.695837

**Authors:** Damla Kaptan, Anne Cecilie Flemming Elvers, Anna Kjær Knudsen, Hannes Schroeder, Hege Ingjerd Hollund

## Abstract

Buried archaeological bones tend to be heavily degraded by microorganisms. This type of biodegradation was already identified in the 19^th^ century and remains a subject of continuous investigation. Yet the specific processes are still not fully understood, and the specific organisms responsible for the decay have not been identified. Technological advances in genetic sequencing now allow detailed study of the bone microbiome. And yet, identifying the species causing the observed bioerosion has proven challenging. Few studies have combined the investigation of bone degradation by microscopy, so-called histotaphonomy, with metagenomic analyses. This study aims to bridge this gap. We utilize a large a set of human bone samples from medieval cemeteries in south-western Norway. Detailed microscopic analyses have been carried out, showing diverse levels of preservation. The extent of bioerosion is correlated with the results from metagenomic analyses as well as environmental factors. Microbiome diversity is greater and more evenly distributed in well-preserved bones with limited bioerosion, particularly those recovered from burials beneath church floors, contrasting with outdoor cemeteries. Our findings show that preservation state is strongly associated with microbiome composition. The most prevalent genus found was *Streptomyces,* supporting previous research suggesting that bacteria within this group could be involved in bone bioerosion.

## Introduction

Bone degradation by microorganisms occurs through enzymatic activities, a process known as bioerosion. After decades of research into bone diagenesis, the organisms causing deterioration of bone remain elusive. The result of the deterioration, on the other hand, is well studied. Microscopy [1,2], porosity measures [3–7], and chemical analyses [8,9] show how microbial action alter the structure and chemistry of buried bones. Understanding bioerosion is crucial for comprehending bone decay due to microbial activity. Already in Child’s article on bioerosion in archaeological bones from 1995, micro-organisms likely involved in bone decomposition were suggested based partially on the characteristics they are expected to have, and partially on cultivation experiments. However, his statement, that ‘…the organisms involved, have not yet been comprehensively defined’ [10] still rings true today. Child (1995) also suggested that work on identification of the organisms involved should be combined with histological analyses of the bone samples [10]. Despite the recent interest in metagenomic analyses and environmental DNA, histological investigation of bone has rarely been combined with analyses of the bone microbiome [11–16]. This study aims to bridge this knowledge gap by investigating how the bone bioerosion and microbiome are correlated, and how the microbiome varies based on the preservation status and the environmental conditions of the biological remains. The ultimate goal is to provide new insights into the types of microorganisms that may be involved in the bone decay process. To address these questions, we analyzed bone samples from medieval cemeteries on the south-west coast of Norway, using light microscopy and scanning electron microscopy to describe and score the bioerosion patterns, and applied metagenome analysis to identify microorganisms. To our knowledge, this is the first time such detailed histotaphonomic analysis of bioeroded bone samples are combined with metagenomic analyses of the bone microbiome.

Bioerosion of skeletal remains has been known since the 19^th^ century, when Wedl made his first observations of tunnels in teeth and bone [17]. Wedl and later scholars have suggested that these were caused by fungi [18,19], however this has been contested and there is still debate on whether some of the tunnelling observed in bone can be connected to fungal activity or not [6,20,21]. The spongiform destructive foci frequently observed in archaeological bones is generally agreed to be caused by bacteria, being made up of remineralized bone mineral and fine sub-micron sized tunnels [6,20,22–24]. Identifying the microorganisms doing the actual bone bioeroding has proven challenging, however.

To be able to directly connect observed bioerosion in bone with the bone microbiome of the exact same sample, we combined microscopic analyses with metagenomic profiling of 83 medieval to post-medieval human bone samples of variable preservation level and assessed the result in relation to environmental conditions.

## Methods and Materials

### Samples and sites

We analyzed bone samples from 83 individuals recovered from six cemeteries along the south-west coast of Norway (Fig 1), dating from the 11^th-^19^th^ centuries. The majority of samples originate from the medieval town of Stavanger and its cathedral cemetery. Additional samples derive from the parish churches of Sola and Hausken, with a smaller number from Ogna parish church, Utstein Monastery, and the royal site at Avaldsnes. The material was primarily obtained through rescue excavations and chance finds from the 1930s to 2022. In addition, 23 of the sampled individuals were uncovered during a research excavation in the old cemetery of Stavanger Cathedral in 2023.

**Fig 1.**
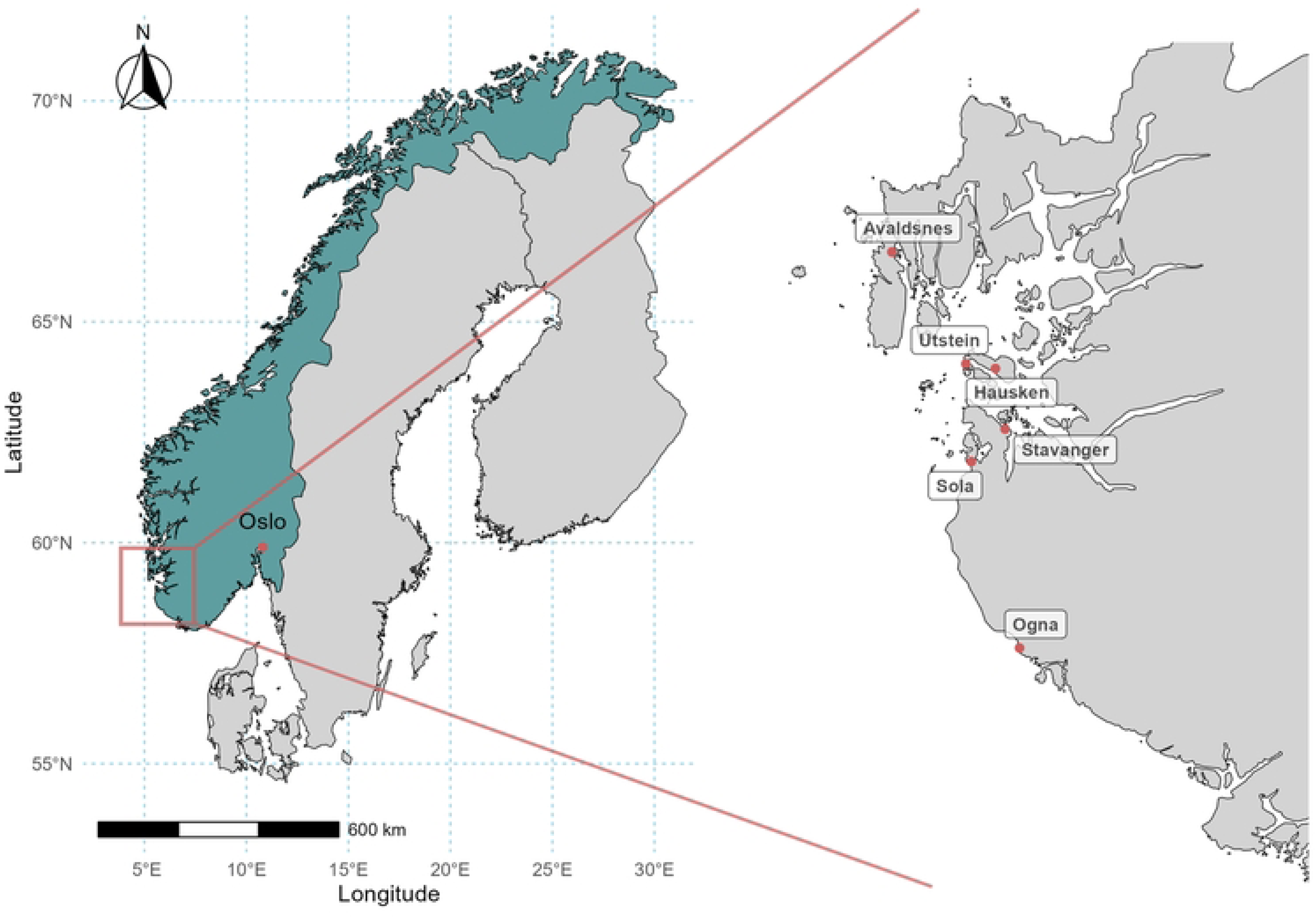
Map of excavation sites.

Due to its robusticity and frequency, the femur is a commonly sampled skeletal element in bioarchaeology. In the current study, this was done when possible; when not possible, the largest available long bone was selected. In instances where only skulls were present, cranial fragments were used (See Table 1, S1 Table and S1 Appendix for more detailed information).

**Table 1.** Site and sample information summarised. Abbreviations for period: *MA* = medieval, *E*= early, *P* = post, *H* = high, *L* = Late, *VA* = Viking Age.

| *Site* | *Excavation year* | *Period* | *N* | *Environment/location* |
| --- | --- | --- | --- | --- |
| *Stavanger East* | 2022 | EMA | 1 | Outdoor |
| *Sola* | 1986 | EMA-PMA | 17 | Outdoor |
| *Ognå* | 1994 | PMA | 3 | Outdoor |
| *Stavanger Byparken* | 2005 | PMA | 4 | Outdoor |
| *Avaldsnes* | 2020 | HMA/LMA | 2 | Outdoor |
| *Stavanger Cathedral nave* | 2021 | LMA-PMA | 12 | Indoor |
| *Hausken* | 2008 | LMA-PMA | 9 | Outdoor |
| *Stavanger Cathedral choir* | 1967 | VA/EMA | 9 | Outdoor |
| *Stavanger Cathedral North* | 2023 | VA/EMA - PMA | 20 | Outdoor |
| *Stavanger Cathedral North* | 2023 | PMA | 3 | Reburied |
| *Utstein* | 1930s-60s | HMA-PMA | 3 | Indoor |

### Histological analysis: bone bioerosion

Transverse sections (∼5 mm) were cut from long bones using a Dremel rotary saw with a diamond blade. Samples were embedded in epoxy resin (Araldite 2020) under vacuum, polished to ∼40 µm, and mounted on microscope slides without coverslips. Thin sections were examined using light microscopy (Olympus BX51) in transmitted normal and polarized light. A subset of samples was further analyzed using a scanning electron microscope (Jeol JSM-IT800) equipped with Oxford Ultim Max 65 mm² EDS, operated either in low vacuum mode (no coating) or high vacuum with silver coating.

Bone preservation was quantified using the Oxford Histological Index (OHI) [25,26], which semi-quantifies bioerosion patterns (Table 2). OHI values (S1 Table) were analyzed in relation to temporal (period), spatial (site and indoor/outdoor), and excavation-year contexts. Multiple linear regression models were fitted using lm() in R to assess the combined effects of sample location (indoor vs. outdoor) and age (recent vs. old) on OHI values. Model diagnostics were examined to ensure validity, and significance was evaluated using ANOVA.

**Table 2.** Oxford Histological Index used for scoring the extent of bioerosion observed in a sample cross-section, in normal transmitted light microscope, following Millard (2001).

| Oxford Histological Index OHI | Approximate % of intact bone |
| --- | --- |
| 0 | <5 |
| 1 | <15 |
| 2 | <50 |
| 3 | >50 |
| 4 | >85 |
| 5 | >95 |

### Ancient DNA extraction and library preparation

Bone samples (S1Table) were manually drilled, and ancient DNA (aDNA) extractions were performed in dedicated cleanroom facilities at the GeoGenetics Sequencing Core, University of Copenhagen.

Briefly, powdered bone samples underwent a short pre-digestion in 1 mL of incomplete digestion buffer (0.5 M EDTA, 30% N-Lauroylsarcosine) at 37 °C for 30 min. The buffer was then replaced with 1.8 mL of freshly prepared digestion buffer (0.5 M EDTA, Proteinase K, and 30% N-Lauroylsarcosine), followed by incubation at 37 °C for 48–72 h on a rocking platform.

aDNA extraction was performed using a protocol adapted for the Biomek i5 Automated Workstation (Beckman Coulter) [27], with the following modifications: For each sample, 150 µL of demineralised lysate were mixed with 1560 µL of binding buffer and 10 µL of magnetic silica beads, and incubated for 15 min with tip-mixing every 5 min. Pelleted beads were washed twice with 450 µL and 100 µL of 80% ethanol + 20% 10 mM Tris-HCl, respectively, and DNA was eluted in 35 µL of 10 mM Tris-HCl + 0.05% Tween-20.

Double-stranded libraries were prepared [28], adapted to the Biomek i5 Automated Workstation (Beckman Coulter), and sequenced on an Illumina NovaSeq 6000 platform at the GeoGenetics Sequencing Core [29].

### Sequence processing and mapping

Raw sequencing files were demultiplexed, and standard Illumina adapter sequences were trimmed using AdapterRemoval v2.3.3 [30], requiring a minimum overlap of 10 bp between read pairs.

### Taxonomic classification and authentication

Taxonomic classification was performed using aMeta [31], a pipeline designed for ancient metagenomic analysis that integrates *k-mer*–based classification via KrakenUniq [32] with competitive Lowest-Common-Ancestor alignment using MALT [33], including additional authentication and validation filters. For KrakenUniq, we employed an extended NCBI non-redundant (NT) database comprising microbial genomes (bacteria, viruses, archaea, fungi, and parasitic worms), the human genome, and complete eukaryotic genomes available at NCBI (downloaded from: [https://scilifelab.figshare.com/articles/online_resource/KrakenUniq_database_based_on_full_NCBI_NT_from_December_2020/20205504]). Reads were filtered to retain only those with at least 1,000 unique *k-mers* and 200 taxonomic reads. Bacteria with taxonomic reads exceeding 500 were processed further with MALT to assess coverage, post-mortem damage (PMD), and edit distance statistics.

### Microbial diversity and statistical analyses

Alpha diversity was calculated using the Shannon index (*diversity()* function, vegan package), with group comparisons performed using Kruskal-Wallis tests and Dunn’s post hoc tests (FSA package, Bonferroni-adjusted). Bray-Curtis dissimilarities were calculated from relative abundance matrices (*vegdist()*), and beta diversity was visualized via *PCoA (cmdscale())* using *ggplot2*.

PERMANOVA analyses (*adonis2()* in vegan package) quantified the influence of environmental factors (age, location, preservation) on microbial composition. Homogeneity of dispersion was tested using *betadisper()* followed by ANOVA to ensure observed differences were due to composition rather than group variance.

All analyses were conducted in R (v4.3.2) within RStudio (2025.05.1 Build 513).

## Results

### Histological analyses: bone bioerosion

Out of the collection of 83 bone samples, 70% displayed bioerosion with an OHI of 4 or less, and 43% were heavily bioeroded with an OHI of 0 or 1. This means that 30% of the samples showed no bioerosion or such limited bioerosion that it did not affect the OHI. Approximately 30% of the entire assemblage displayed intermediate levels of bioerosion, where larger parts of the bone remain unchanged (OHI of 2-4) (S1Table). The well-preserved areas of bone in all samples, including the ones with OHI of 5, exhibited enlarged osteocyte lacunae and canaliculi, and diagenetic pores that had not yet developed into the classic destructive foci often observed in archaeological bone. These are possibly early-stage microbial decay [34]. This suggests that bioerosion was initiated in most instances, but further decay was inhibited or slowed down (Fig 2).

**Fig. 2.**
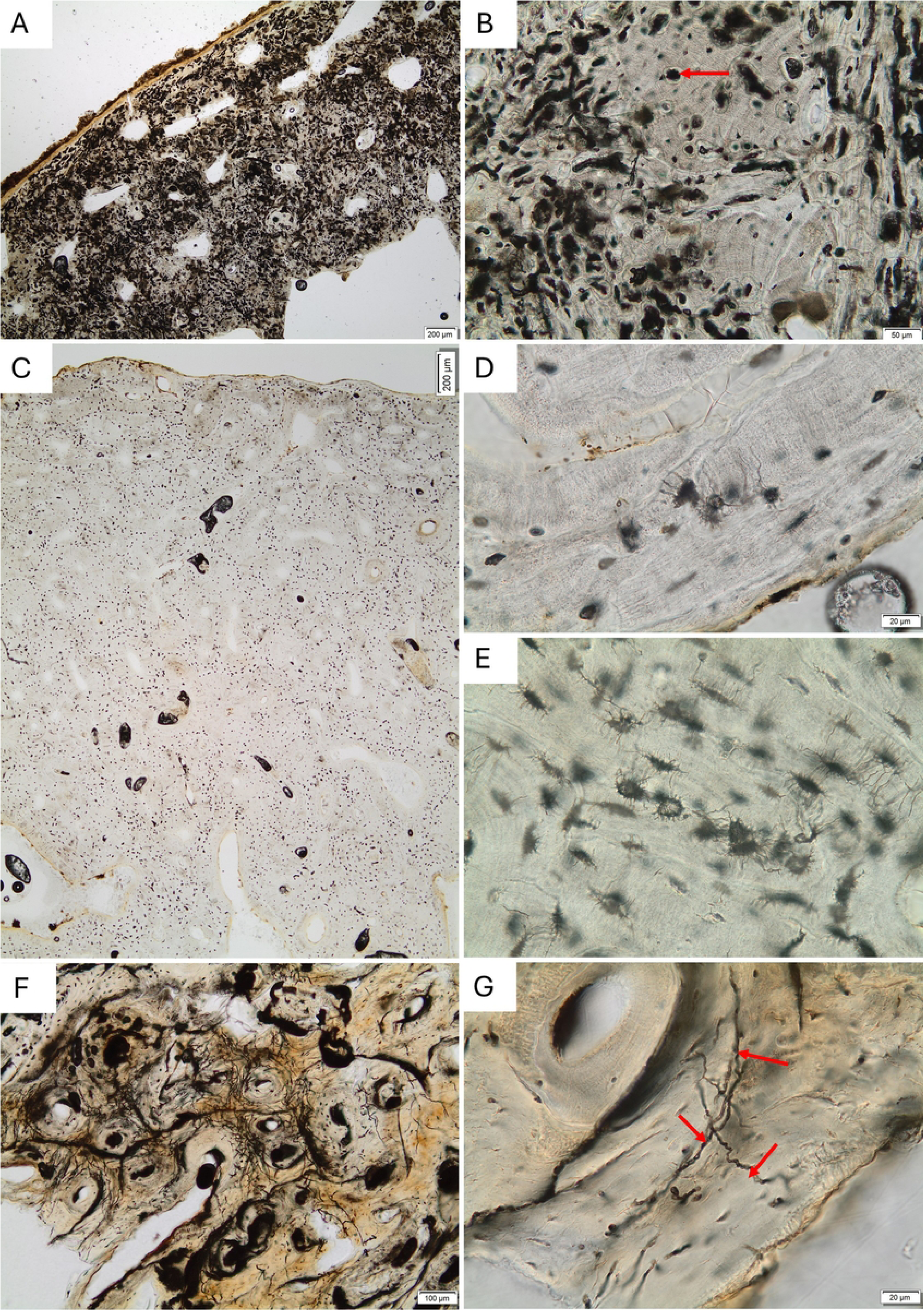
Examples of bioerosion in the studied assemblage. (A) Extensive bioerosion in sample S10294.17b; no original microstructure is visible. (B) Detail from the same sample as in (A), showing characteristic destructive foci and small areas of well-preserved bone despite intense microbial attack. The arrow indicates a destructive focus surrounded by preserved bone. (C) Well-preserved sample S14393.104 with no destructive foci observed. (D) Detail of the same sample, showing enlarged osteocyte lacunae and canaliculi. (E) Similar features as in (D) observed in another sample, S14393.80. (F–G) Wedl-type tunnelling observed in the mid-cortex of sample S14604.10 at two magnifications. Arrows in (F) indicate individual tunnels.

Fungal remains were observed in many thin sections, in the form of hyphae, fungal fruiting bodies, and spores. The fungal remains were mainly found within cracks and natural pores in the bone on or close to bone surfaces (S1 Fig). Two different types of hyphae were observed, often in the same sample, with different thickness, one roughly 1-2 microns across, and the other 5-10 microns across. Both were reddish-brown color. The thicker hyphae were clearly branching; the thinner hyphae were sometimes collected in dense bundles of hyphae. Different shapes of fungal spores were also observed (S1 Fig).

Possible biofilm was also observed in the microscope, but the nature and extent of this is not possible to identify and quantify based on visual examination in the microscope. Light and electron images of examples can be seen in S2 Fig.

Most samples with little or no bioerosion were from burials inside still standing buildings, such as the Utstein Monastery church, and Stavanger Cathedral nave, whereas bioerosion was common in the outdoor burial grounds. However, bioerosion was also observed in a few samples from indoor burials, and there was a great variation in OHI scores at the outdoor cemeteries of Hausken and Stavanger (S2 Fig).

During the 2023 research excavation in Stavanger, three phases and groups of skeletal assemblages were apparent: 1) *In-situ* post-medieval burials (level I), 2) *in-situ* medieval burials (level II), and 3) re-deposited post-medieval coffins. The re-deposited remains, likely removed from the cathedral’s burial chambers in the 19^th^ century and reburied outside, display no bioerosion. However, only three of the nine individuals uncovered were sampled. Conversely, level II medieval burials were all heavily bioeroded. The *in-situ* post-medieval burials from level I also displayed extensive bioerosion, but more than half (6 out of 11) had an OHI of 2 or higher, and three samples were scored with an OHI of 5.

To assess the potential influence of period, site, and excavation year on bioerosion, we analyzed the OHI scores under varying conditions. An increasing trend in OHI was observed in more recent samples compared to those from older periods (Fig 3A). Further analysis of OHI values across different sites indicated that the highest OHI values were observed in indoor environments (Fig 3B). By categorizing the samples into “Recent” (post-medieval periods) and “Old” (earlier than post-medieval), a significant difference in the distribution of OHI values between recent and old samples was evident (Fig 3C). A detailed comparison between indoor and outdoor environments showed a significant difference, with indoor sites exhibiting higher OHI values (Fig 3D). However, the comparison of excavation year and OHI values did not reveal any specific relationship, apart from the observed pattern influenced by the indoor/outdoor effect (S3 Fig).

**Fig 3.**
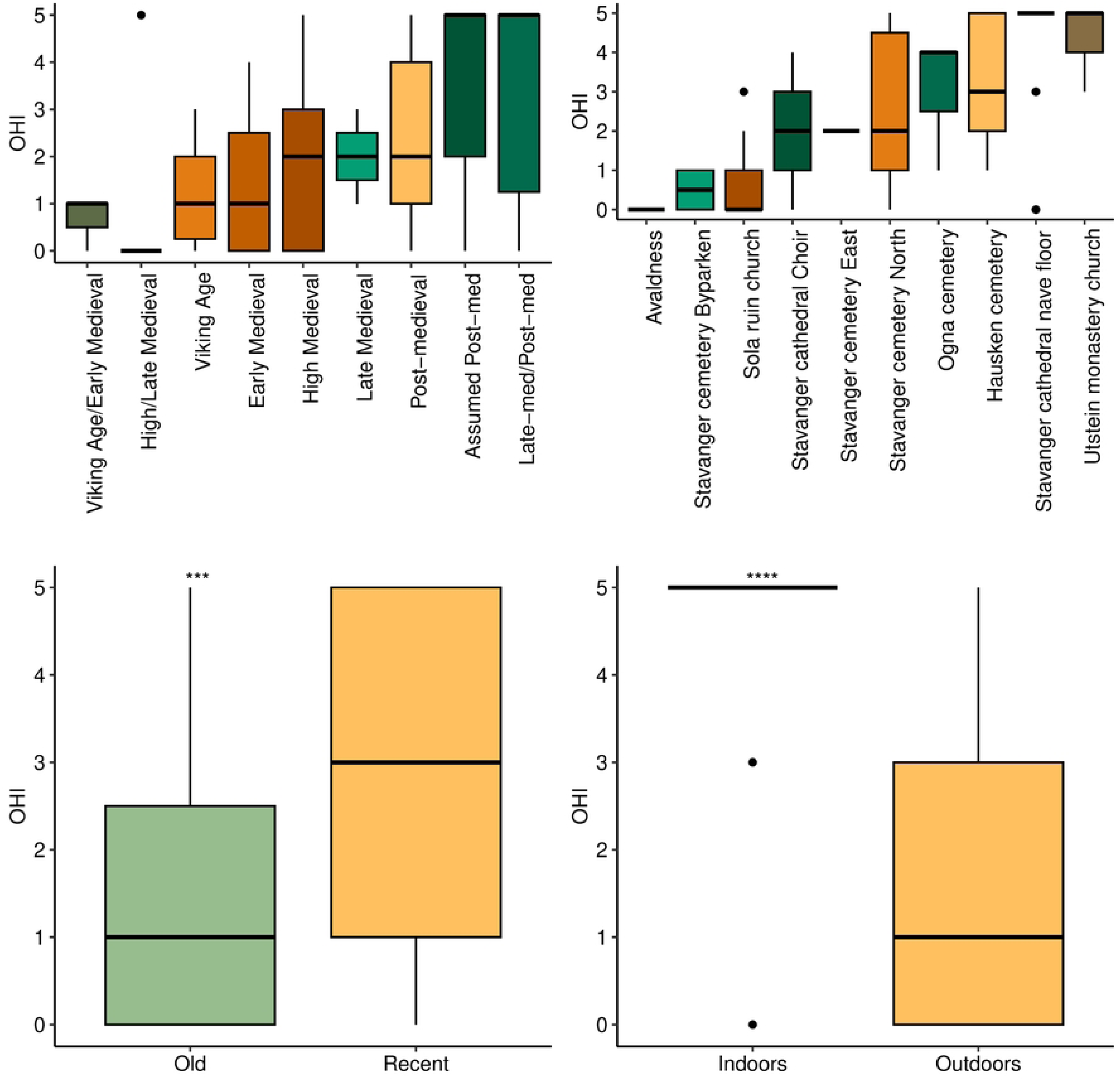
Correlation between OHI values and different parameters. (A) Sample period. (B) Excavation site. (C) Sample age (recent includes all post-medieval samples; old includes all earlier periods). (D) Sample location. Statistical significance was assessed using the Wilcoxon rank-sum test (*** p < 0.001; **** p < 0.0001).

A multiple linear regression analysis was conducted to examine the effects of location (indoor vs. outdoor) and age (recent vs. old) on OHI values. The regression model was statistically significant, (F(2, 80) = 22.39, p<0.001) and explained approximately 35% of the variance in OHI values (R^2^adj=0.3428). Specifically, the coefficient for outdoors samples was -2.28 (p<0.001), indicating that outdoor samples were associated with significantly lower OHI values compared to indoor samples. Additionally, the coefficient for recent samples was 1.29 (p<0.001), suggesting that more recent samples had significantly higher OHI values compared to older samples. The residuals of the model were approximately normally distributed, with a residual standard error of 1.6 on 80 degrees of freedom. The minimum and maximum residuals were -4.78 and 2.79, respectively.

### Metagenomic analyses: the bone microbiome

Extraction and library blanks, processed identically to the samples, showed no detectable microorganisms. KrakenUniq classification, however, identified 29 bacterial species and one fungal species spanning 16 genera in 83 individuals (S2 Table). Microorganisms exceeding 500 taxonomic reads in KrakenUniq were further evaluated using MALT. Only four species had more than 5,000 reads, each with a breadth of coverage exceeding 5% and an average depth of 0.08× (S3 Table), with postmortem damage (PMD) profiles indicative of ancient origin. To enhance the reliability of taxonomic assignments, we restricted downstream analyses to the genus level and excluded any genera present in only a single individual, resulting in 13 bacterial genera retained across all samples. The most prevalent genus was *Streptomyces*, detected in 86% of individuals (Fig 4), followed by *Streptosporangium* and *Lysobacter*. In addition to bacterial genera, the fungal genus *Serpula* was detected in one sample from Stavanger Cathedral nave (S14393.72).

**Fig 4.**
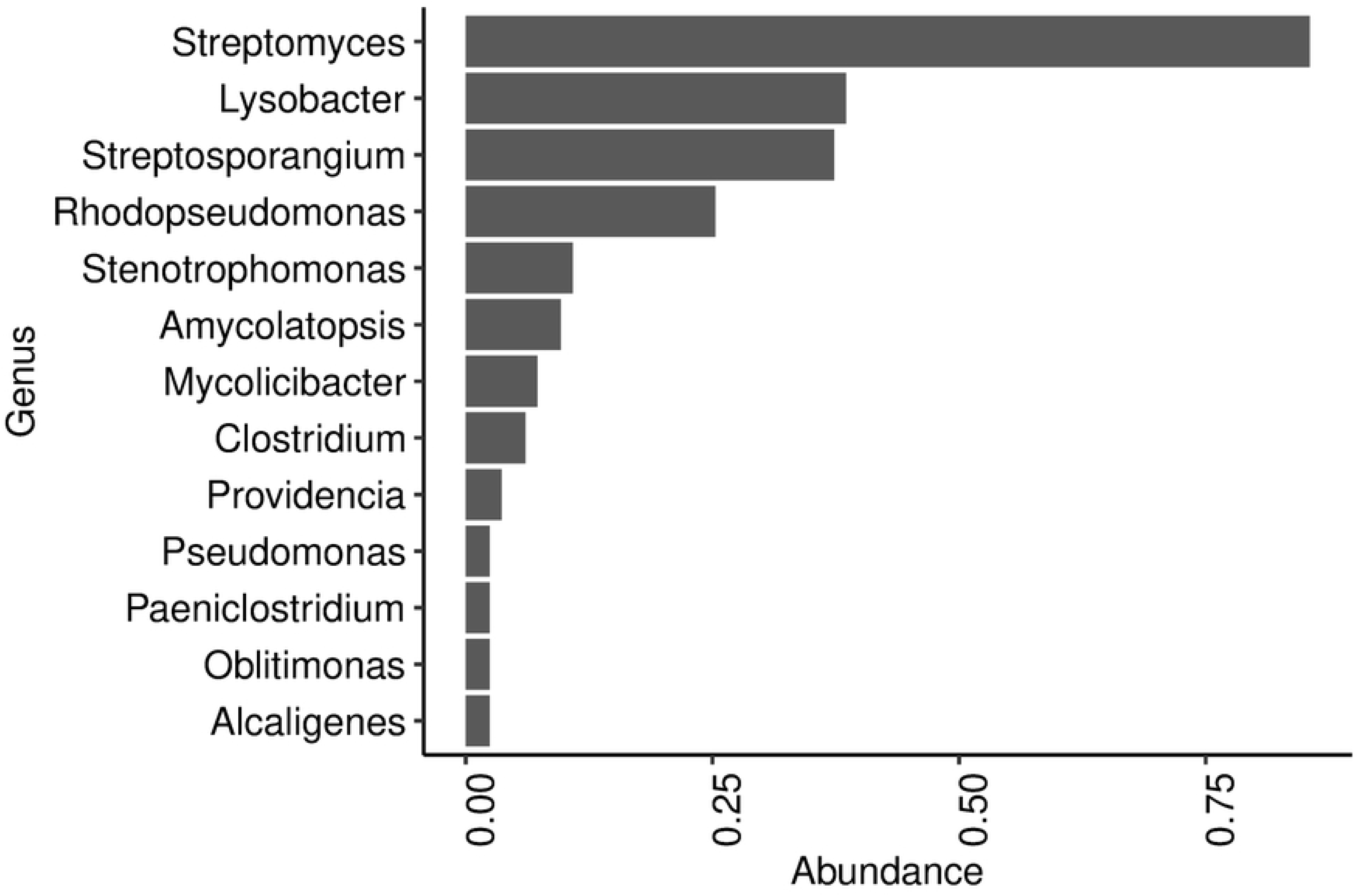
Microbiome composition across all samples. Only microbial genera observed in at least two samples are plotted.

Microbial alpha diversity, assessed using the Shannon index, was lower in “old” and/or “outdoor” samples compared to “recent” and/or “indoor” ones (Wilcoxon rank-sum test, *p* < 0.005; S4A–B Fig). Similarly, Bray–Curtis dissimilarity analyses revealed significant effects of both sample age (*p* = 0.03) and storage location (*p* = 0.003) on microbiome composition (Kruskal–Wallis test; S4C–D Fig). No significant correlation was observed between excavation year and microbial diversity (S5 Fig).

To further investigate potential storage effects, we compared well-preserved outdoor samples (OHI 4–5) grouped by excavation period: *old excavations* (Stavanger 1967, Ogna 1994, and Hausken 2008) and *recent excavations* (Stavanger 2023). Although museum-stored samples from older excavations tended to show slightly higher microbial diversity than freshly excavated ones, the difference was not statistically significant (S6 Fig).

PERMANOVA analyses on the full Bray–Curtis dissimilarity matrix confirmed significant effects of age (F = 3.23, R² = 0.048, *p* = 0.035) and location (F = 6.95, R² = 0.10, *p* = 0.001). A combined model including both factors explained 14% of the variance (F = 4.99, R² = 0.14, *p* = 0.001). The assumption of homogeneity of group dispersion was met for both *Age* (F = 1.81, *p* = 0.18) and *Location* (F = 0.09, *p* = 0.77), indicating that the observed differences in microbiome composition are not attributable to unequal within-group variance.

Observing that both the OHI and microbiome composition are influenced by environmental conditions, we investigated whether their correlation supports this pattern by assessing the impact of bioerosion on microbiome abundance. Samples were grouped by OHI scores, and genus-level microbial composition was visualized (Fig 5). In samples with extensive bioerosion (OHI = 0, 1), *Streptosporangium* was the most dominant genus. In samples with moderate bioerosion (OHI = 3,4), *Lysobacter* predominated, while in well-preserved samples (OHI = 5), *Streptomyces* was the most abundant genus. Notably, samples with minimal bioerosion (OHI = 5) exhibited greater genus-level diversity compared to more degraded samples (OHI < 5).

**Fig 5.**
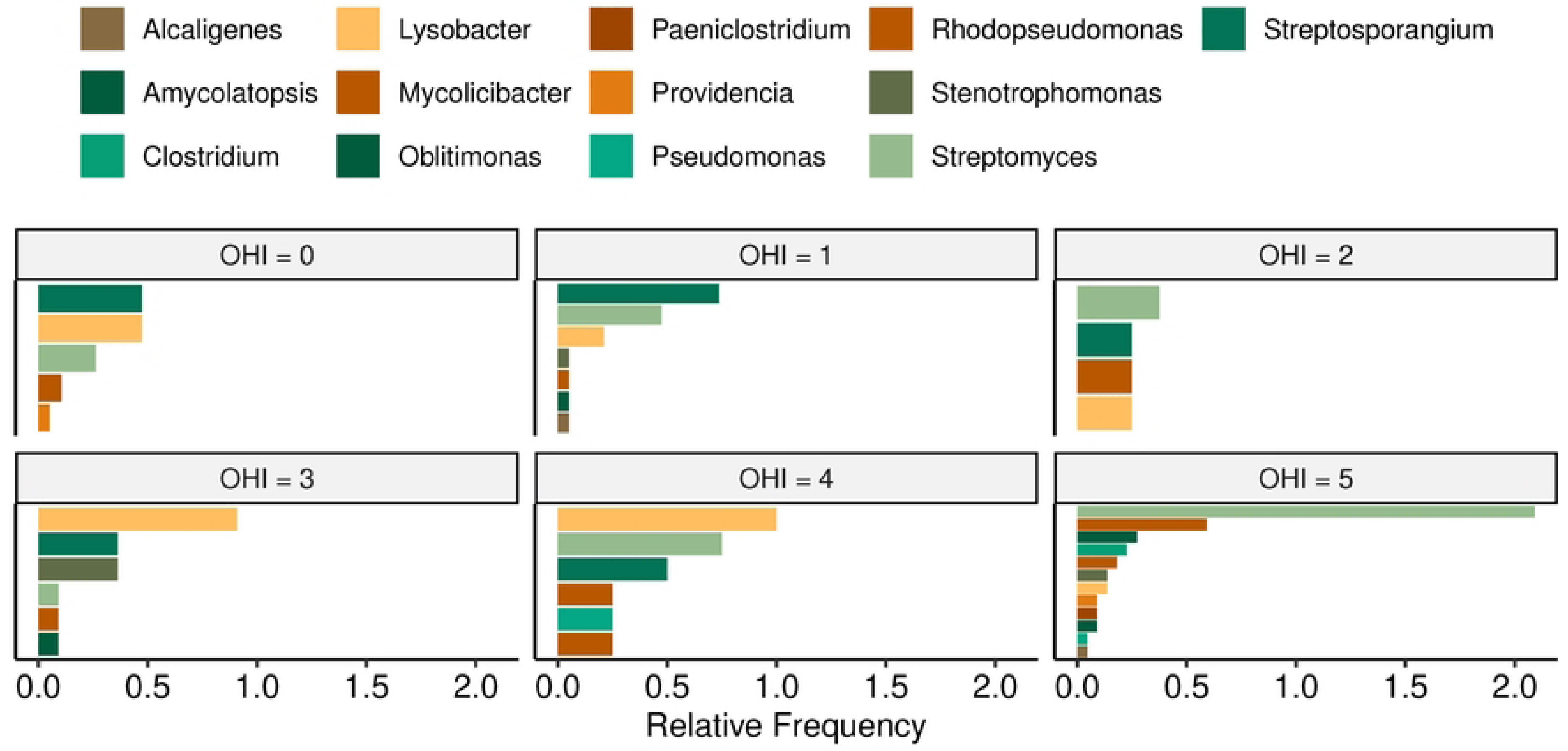
Microbial composition of samples with different OHI scores.

To assess within-sample microbial diversity, Shannon diversity indices were calculated across samples grouped by preservation state (OHI levels) (S7A Fig). A Kruskal-Wallis test revealed a significant effect of preservation state on Shannon diversity (χ² = 15, df = 5, *p* = 0.01). Pairwise comparisons using Dunn’s test (Bonferroni-adjusted) showed that samples with the highest preservation level (OHI 5) had significantly higher diversity than OHI 0 and OHI 1 samples (*p* = 0.05, and 0.03 respectively).

PERMANOVA analysis based on Bray–Curtis dissimilarities indicated a significant effect of preservation state on microbiome composition (F = 17.7, R² = 0.22, *p* = 0.001), suggesting that the evenness of microbial communities differed across preservation categories. A test for homogeneity of multivariate dispersion showed no significant differences among categories (PERMDISP: F = 2.09, *p* = 0.079), indicating that the PERMANOVA result was not confounded by unequal within-group dispersion. To further explore these differences, the distribution of the first principal coordinate (PCoA1) from Bray-Curtis distances was examined. PCoA1 values varied significantly among OHI groups (Kruskal-Wallis χ² = 23.8, df = 5, *p* = 0.0002) (S7B Fig). Post hoc Dunn tests identified significant differences mainly between the lowest preservation states (OHI 0 and 1) and the highest (OHI 5), indicating that better-preserved samples harbor distinct microbial communities. Overall, these results show that well-preserved bones tend to support richer and more evenly distributed microbial communities.

To visualize microbiome patterns across sample sets varying in location and age, we compared microbial abundances between indoor nave burials and outdoor North burials at Stavanger Cathedral. The North burials were further subdivided into: North Level I (post-medieval, similar to the nave), North Level II (Medieval), and North redeposited burials (originally interred in 17^th^ century grave chambers indoors and moved to the North graveyard in the 19^th^ century). Indoor (nave) samples displayed a richer and more diverse microbiome, in contrast to North Level II burials, which showed only *Streptosporangium*. North Level I samples had more diversity than Level II but remained poorer than the Nave samples, while redeposited burials displayed an intermediate pattern between indoor and outdoor groups (Fig 6). These microbiome differences closely mirrored the OHI distribution, with the Nave exhibiting the highest preservation, North Level II the lowest, and a more diverse preservation pattern in the rest. Together, these findings demonstrate that preservation state is strongly associated with microbiome composition, reflecting and interacting with burial environment and postmortem conditions.

**Fig 6.**
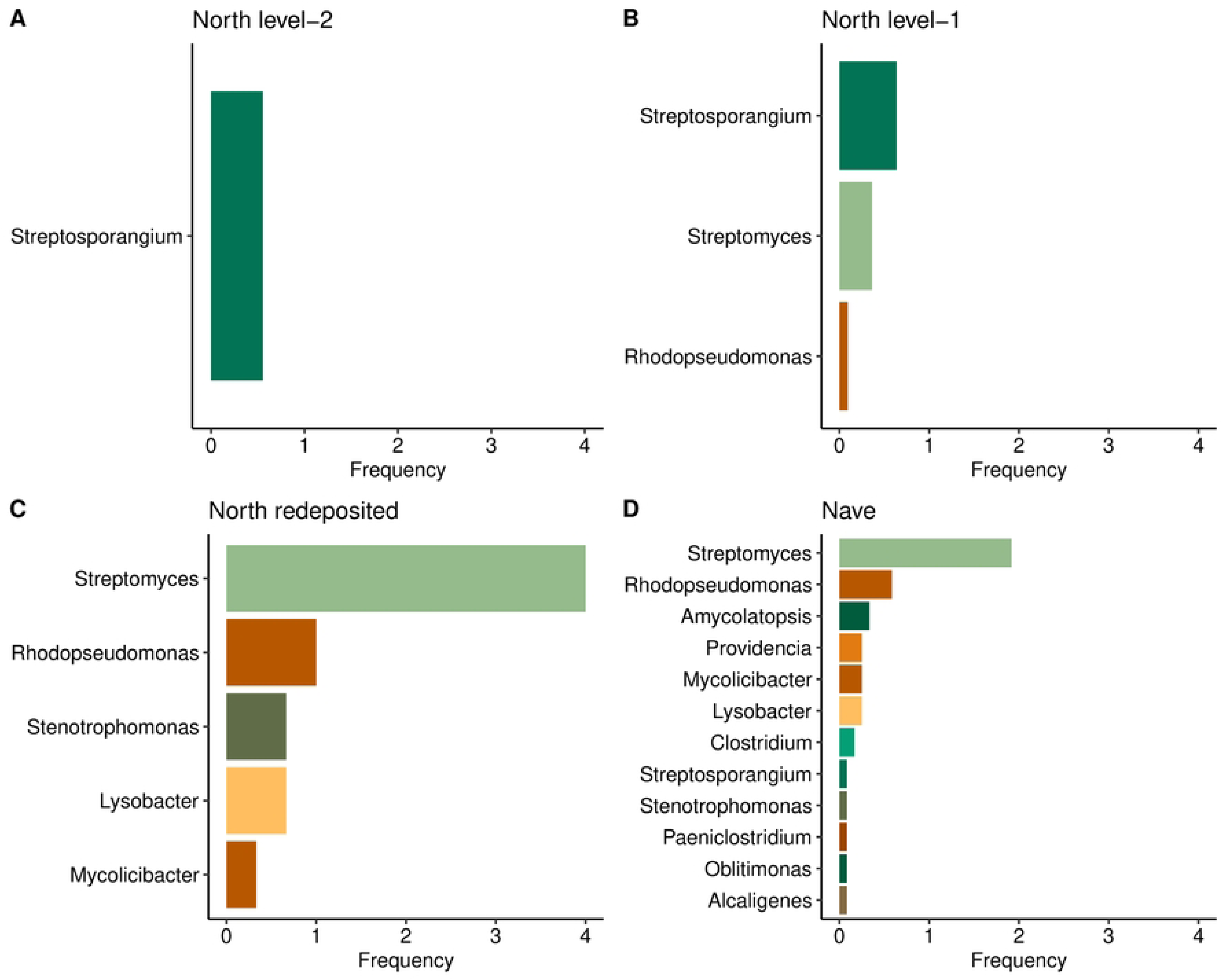
Relative abundances of microbial genera across different burial contexts. (A) North level 2. (B) North level 1. (C) North redeposited, and (D) Nave burials. For each genus, presence/absence across samples was recorded, and the total presence count was normalized by the number of samples in each group.

## Discussion

In this study, we utilized a large set of samples displaying different preservation levels, and combined histotaphonomic and metagenomic investigations. This diverse collection provides an ideal framework for exploring the variation in the metagenome, based on the presence, absence, and extent of bioerosion. The study represents a detailed analysis of human burials, encompassing various burial sites, periods, and excavation years, utilizing metagenomics. The aim is to understand bone histology and bioerosion in comparison with the metabiome.

### Environmental impacts on bone bioerosion and metagenome

In this study, our primary interest lies in the degradation of biological remains in various burial environments. However, due to the inability to conduct environmental screening and control collection for older excavations, we lack the data required for an in-depth analysis of environmental factors affecting bioerosion. Nonetheless, the basic information obtained indicates that more recent biological material is better preserved in indoor burial environments, which is valuable for future conservation efforts.

Regression analyses identified burial location and age as significant predictors of histological preservation (OHI), explaining ∼34% of variance. Both variables also influenced microbial diversity, confirmed by PERMANOVA, indicating that temporal and environmental factors independently shape bone microbiomes. Recent indoor samples displayed richer, more even microbial communities than older outdoor ones, despite the latter often showing greater bioerosion.

Both bone and the bone metagenome are generally better preserved in younger burials within an indoor environment, compared to old burials and/or outdoor cemeteries. Notably, recent samples from indoor contexts harbor richer and more evenly distributed microbial communities than older samples from outdoor burials. Despite the detection of diverse microbial taxa in the bones, the indoor conditions do not appear conducive to microbial degradation of bone. This is according to expectation, as the indoor materials are shielded from percolating rainwater and fluctuating burial conditions. Indeed, environmental monitoring of the cultural layers beneath the cathedral floor has shown that the sediments are relatively dry and exhibit a weakly alkaline pH [35], which is beneficial for the preservation of both bone [22,36,37] and DNA [38–40]. The alkaline pH, along with low moisture and minimal water movement, likely limited microbial access to bone proteins by preventing mineral dissolution, explaining the low bioerosion [20,41]. The preservation of wood, textiles, and plant remains in some graves [42], further attests to the protective nature of the indoor burial environment. Other environmental factors—such as infiltration of humic substances, organic decomposition byproducts, and metal ions from corroding artifacts [43,44] —may have rendered the bone matrix less susceptible to microbial activity. In contrast to the outdoor environment where only iron rivets from coffins were preserved alongside the skeletons, numerous metal objects were found beneath the nave floor [42].

Nine individuals in the Stavanger Cathedral North excavation were originally chamber burials from the 17^th^ century, exhumed and reburied outside in the 19^th^ century (S1 Appendix). These represent a unique environmental and taphonomic history. When uncovered, all were in anatomical position, indicating that soft tissues were likely still intact at the time of reburial. Most likely, the chamber conditions were dry, leading to mummification by desiccation as reported in other above-ground 17^th^ century graves [45]. Following reburial in soil, the remains became fully skeletonized, but the level of preservation is quite different from the in-situ burials. In two cases, hair was still preserved, and remnants of wooden coffins were present. Only three out of nine individuals discovered were analyzed. All exhibited an OHI of 5 with no bioerosion. It is likely that the drainage trench into which the coffins were reburied, along with the coffin materials themselves, created a distinct microenvironment, differing from the rest of the excavation area. Comparisons of the microbiomes of these redeposited individuals revealed an intermediate pattern between the microbially diverse indoor (nave) samples and the low-diversity outdoor (north) burials (Fig 6). Further comparisons among the North burial subgroups reinforced this pattern of ongoing decay: North Level II burials, which were deeper and older, exhibited extremely limited microbial diversity, with only a single genus detected, whereas North Level I burials displayed slightly richer microbiomes. The detection of *Streptosporangium* as the sole genus in North Level II burials may indicate a unique group of microorganisms adapted to survive under nutrient-limited conditions in highly bioeroded bones. However, this hypothesis requires confirmation through further research on *Streptosporangium*.

Several sub-collections of the skeletal assemblage under study were excavated decades ago, the earliest in the 1930s (S1 Appendix). It seems that excavated bones contain active microorganisms, and that the profile of the bone microbiome of museum-stored bones is different from the soil and from freshly excavated bones [12,16,46]. In our study, although the material stems from the same region and a relatively narrow time range, we do not have any directly comparable assemblages of bones from the same site but excavated at different times. Furthermore, parts of the collection were stored at different institutions, and we do not have detailed information on the storage environment throughout the history of the collections. Correlating bone preservation and microbiome there is no relationship with the year of excavation and site effects are prevalent (S3 Fig). Heavily bioeroded bones displayed low microbial abundance (see discussion below) regardless of excavation year. However, the pattern seen when well preserved bone (OHI of 4 or 5) from outdoor cemeteries is compared, concurs with the results of the previous studies where museum stored bones display a greater microbial diversity, suggesting that microbial life continues in storage, and that a different microbiome to that of the soil develops [12,16,46].

### Microbial correlates of preservation

The complete aMeta pipeline resulted in only four microorganisms being confidently identified, as most taxa either lacked sufficient numbers of mapped reads or displayed irregular patterns of genome coverage for the species detected by KrakenUniq. A notable example is *Streptosporangium roseum*, which appeared in the majority of samples according to KrakenUniq but showed an inconsistent signal when re-evaluated with MALT. Although KrakenUniq reported numerous reads, MALT mapping revealed that these reads were confined to a few genomic regions and exhibited high mismatch rates, indicating that the assignment to the *S. roseum* reference genome is unreliable. A parsimonious explanation is that the detected reads derive from a related organism lacking a close representative in current reference databases—or possibly from a yet-undescribed taxon—rather than from *S. roseum* itself. Given the ancient and poorly preserved nature of the analyzed material, low read counts were expected. Therefore, to ensure reliability, we report all detected taxa conservatively at the genus level rather than as confirmed species.

Microbiome composition correlated strongly with OHI, with better-preserved bones (OHI 5) showing greater alpha diversity than degraded bones. This likely reflects a continuum in which well-preserved bones retain more nutrients and structural integrity that support colonization, whereas heavily degraded bones lack sufficient substrates. Taxonomic patterns support this interpretation. *Streptomyces*, a common soil genus capable of degrading complex organics [47,48], predominated in well-preserved bones, consistent with findings from Neanderthal and reindeer remains [12,16]. In contrast, *Streptosporangium*, another actinobacterium, was more abundant in heavily bioeroded bones, while *Lysobacter* appeared mainly in moderately degraded ones. Notably, species from both *Streptomyces* and *Streptosporangium* genera encode M09A-type collagenases—enzymes capable of degrading bone [12]. These distribution patterns may reflect stages of microbial succession or preservation-linked ecological niches, although their precise roles in bone diagenesis remain unresolved.

The fungal genus *Serpula* was detected in only one sample in the KrakenUniq analysis. Although intriguing—possibly reflecting specific microenvironmental conditions—insufficient coverage prevents confident interpretation. This finding should therefore be viewed with caution.

Nonetheless, fungal structures were observed histologically in this and several other samples. Wedl-type tunnels, sometimes attributed to fungal activity [6,18,19,21], were also present in some specimens (S1 Fig). These fungal remains were generally confined to the outer bone surfaces, with only a few scattered microscopic fragments within the tissue. Most were likely removed during sample preparation, when outer layers were cleaned prior to DNA extraction. Together, the histological and metagenomic evidence suggests that fungi do not play a central role in the bioerosion of bone and that the observed bioerosive patterns are predominantly bacterial in origin. Importantly, our ancient DNA data were generated as part of another ongoing study primarily focused on bone and DNA preservation. Here, we repurposed the same dataset to examine microorganisms potentially involved in bone diagenesis. Because only ancient DNA library preparation methods were applied, modern DNA and contemporary microorganisms, including recent fungal growth, were likely underrepresented, potentially limiting detection of the extant microbiome.

While our DNA-based analyses provide a comprehensive overview of microbial presence in archaeological bone, future studies incorporating RNA-based metatranscriptomic approaches could help capture the fraction of the microbiome that is potentially metabolically active. Transcriptomic data would allow the identification of genes involved in collagen and mineral degradation, providing functional insight into microbial contributions to bone bioerosion. Combining DNA and RNA profiling would therefore offer a more dynamic view of microbial activity and succession within the bone microenvironment, complementing the preservation and taxonomic patterns observed in this study. Overall, our results indicate that bone microbiomes are shaped by both preservation state and burial environment, and that microbial communities in archaeological bone may actively contribute to degradation rather than merely reflect contamination. The higher microbial diversity observed in well-preserved bones suggests that long-term preservation may coexist with microbial activity, underscoring the complex and dynamic nature of bone diagenesis.

## Conclusion

In this study, we made a direct comparison between the extent of bone bioerosion and composition of the bone microbiome, using a substantial and diverse collection of human bone samples from historic cemeteries and churches in south-western Norway. By analysing correlations between bone preservation, microbial communities, and environmental conditions, we found that the highest microbial diversity was associated with well-preserved bones and indoor burials. Due to the nature of the sampled materials, we were unable to confidently assess the effects of post-excavation storage on microbial composition. Pinpointing the specific organisms responsible for bioerosion remains challenging, particularly given the limited taxonomic resolution of ancient DNA data, which prevented species level identification. However, the results align with previous research suggesting the existence of a distinct “museum bone microbiome”, and highlight the central role of *Streptomyces* bacteria in bone degradation during soil burial. Overall, our results contribute to a more nuanced understanding of bone bioerosion. Future research from other geographical regions and site types, particularly using well-documented and systematically sampled collections, will help to build on these findings and further clarify the mechanisms driving bone degradation.

## Acknowledgements

We are grateful to Bent Philippsen for assistance with accessing and utilizing the servers at the GeoGenetics Sequencing Core. We thank Espen Undheim (University of Stavanger) for support with the SEM analyses, and Tom Gilbert (University of Copenhagen) for guidance regarding the genetics aspect of this study. We also thank Arda Sevkar (Hacettepe University) for support in metagenome analysis and for sharing his Bash scripts on aMeta.

Declaration of generative AI and AI-assisted technologies in the manuscript preparation process

During the preparation of this work, the author(s) used ChatGPT to improve the grammar and clarity of the manuscript. After using this tool/service, the author(s) reviewed and edited the content as needed and take(s) full responsibility for the content of the published article.

## Supporting information

**S1 Fig.**
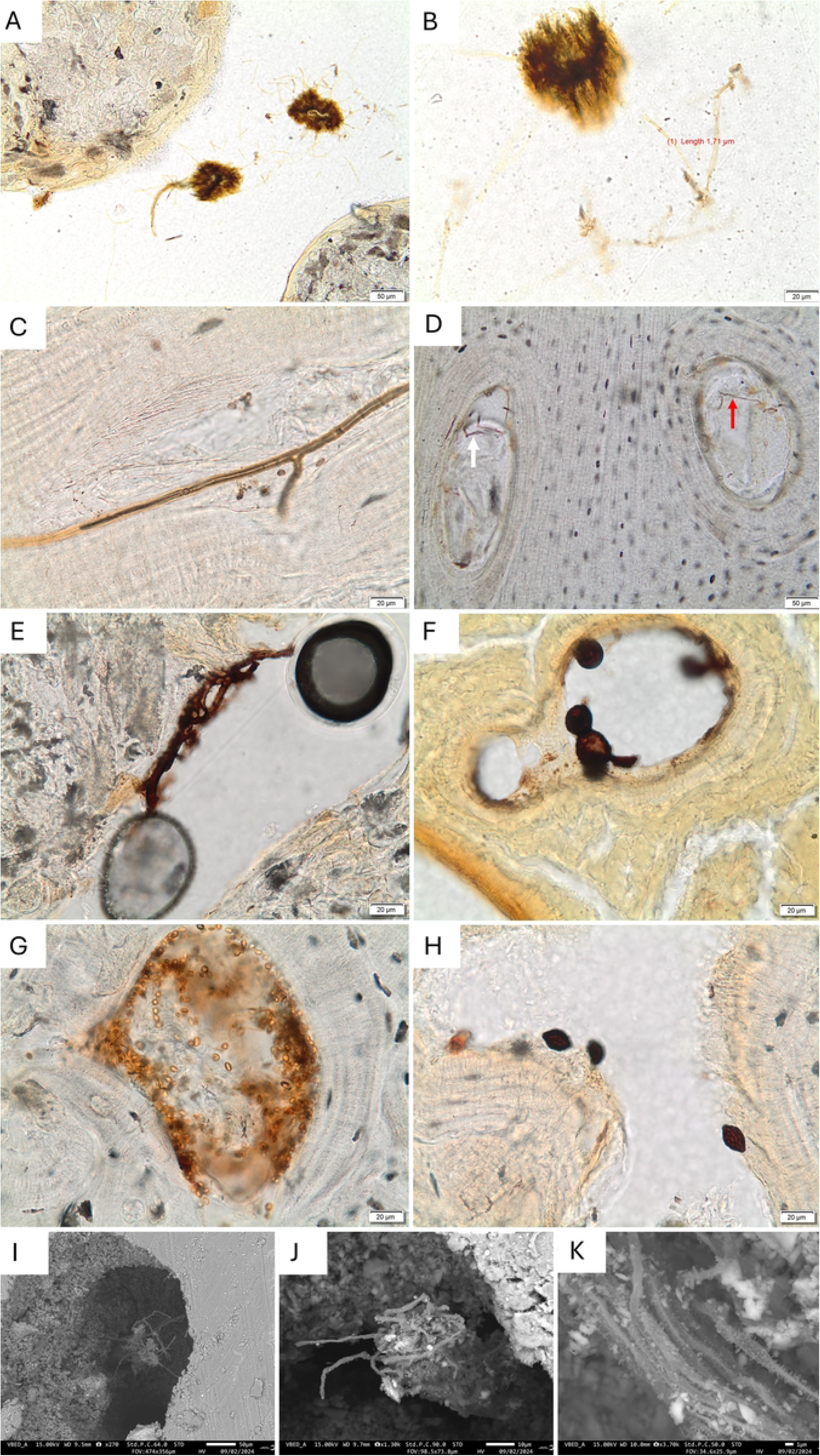
Microscopic and SEM Evidence of Fungal Structures. (A) Fine hyphae and bundles of hyphae within the trabecular bone of S14393.83. (B) Detail of the fine hyphae and bundles in the same sample as in (A). (C) Thicker hyphae in sample S14393.72, clearly branching and segmented. The brown grains seen along the hyphae could be spores, or the cross-section of similar hyphae. (D) Fine hyphae (arrows) within two Haversian canals (blood channels) of sample S14393.118. (E) Fungal hyphae in a Haversian canal in S10294.18. The two large spheres are air bubbles in the embedding resin. (F) Spherical spores or sporangia within a Haversian canal in SZ20610. (G) A mass of almond-shaped spores filling a Haversian canal in SZ21164. (H) Lemon-shaped fungal spores in S14393.72. (I-K) SEM-images in backscatter mode of fine fungal hyphae, in pores of an unpolished bone sample of SZ20839. The diameter is roughly one micron and the hyphae have a hairy surface.

**S2 Fig.**
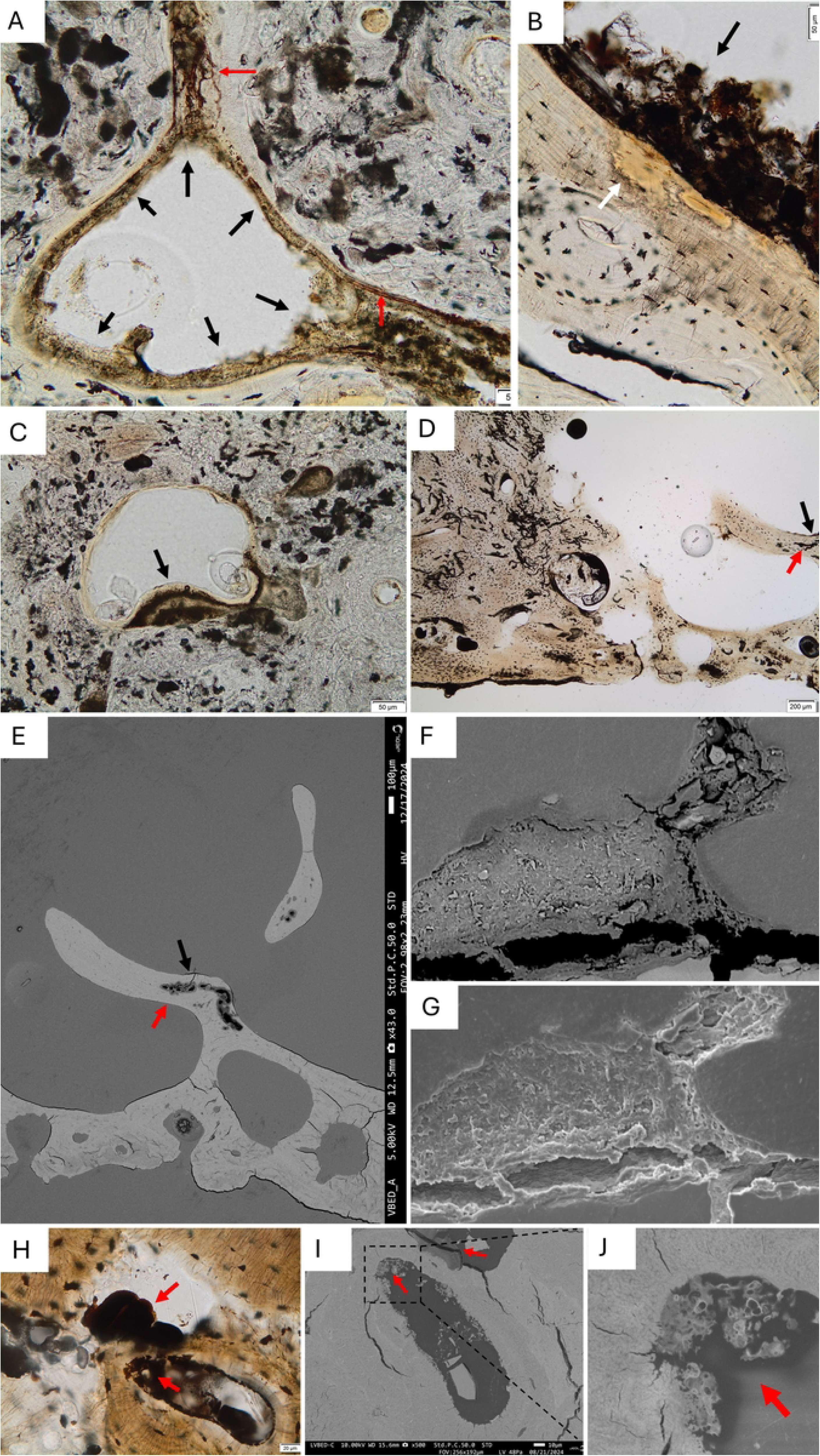
Examples of possible biofilm observed under light and electron microscopy. (A) Fine hyphae and bundles of hyphae within the trabecular bone of S14393.83. (B) Detail of the fine hyphae and bundles in the same sample as in (A). (C) Thicker hyphae in sample S14393.72, clearly branching and segmented. The brown grains seen along the hyphae could be spores, or the cross-section of similar hyphae. (D) Fine hyphae (arrows) within two Haversian canals (blood channels) of sample S14393.118. (E) Fungal hyphae in a Haversian canal in S10294.18. The two large spheres are air bubbles in the embedding resin. (F) Spherical spores or sporangia within a Haversian canal in SZ20610. (G) A mass of almond-shaped spores filling a Haversian canal in SZ21164. (H) Lemon-shaped fungal spores in S14393.72. (I-K) SEM-images in backscatter mode of fine fungal hyphae, in pores of an unpolished bone sample of SZ20839. The diameter is roughly one micron and the hyphae have a hairy surface.

**S3 Figure.**
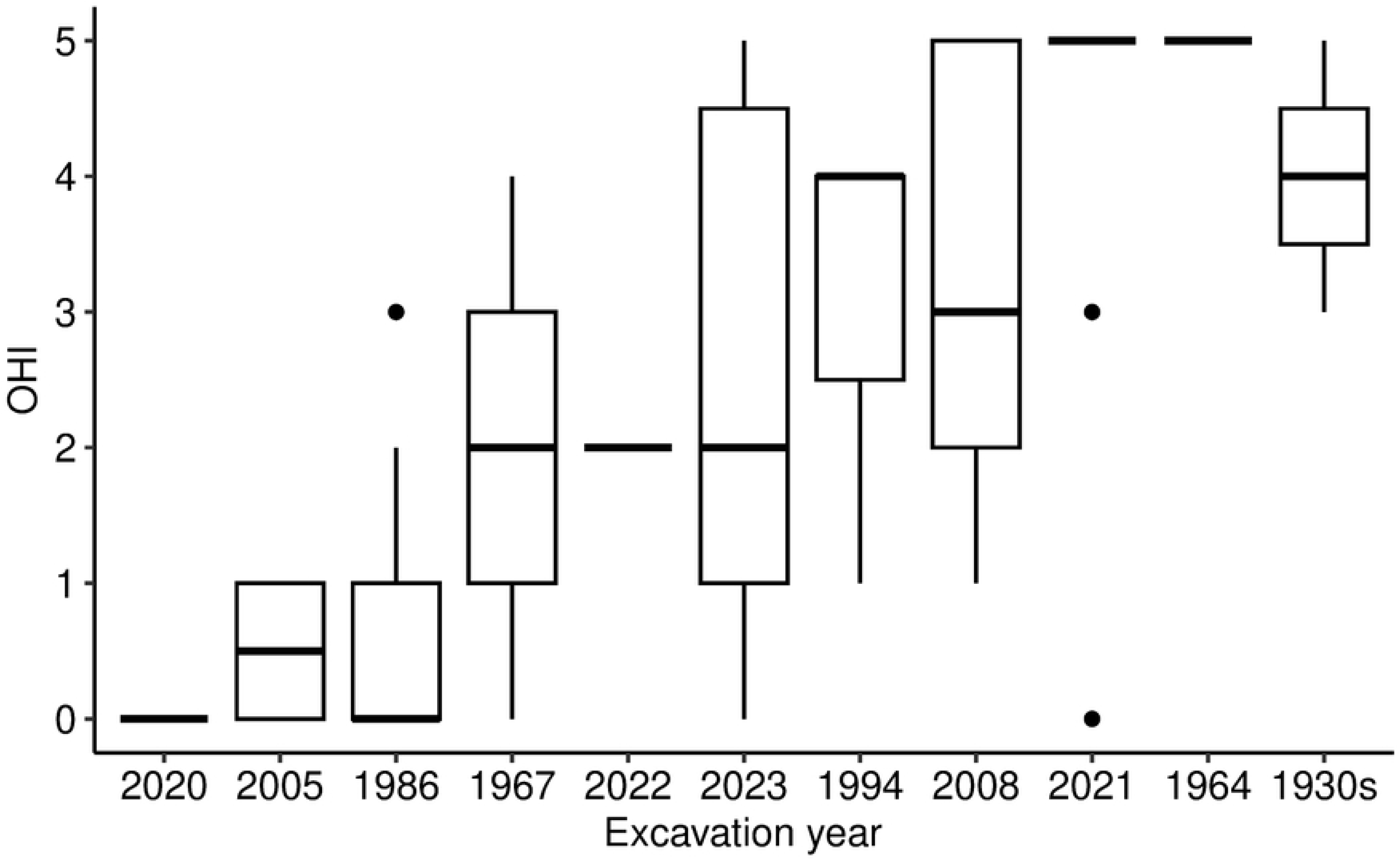
Comparison of OHI values with excavation year.

**S4 Figure.**
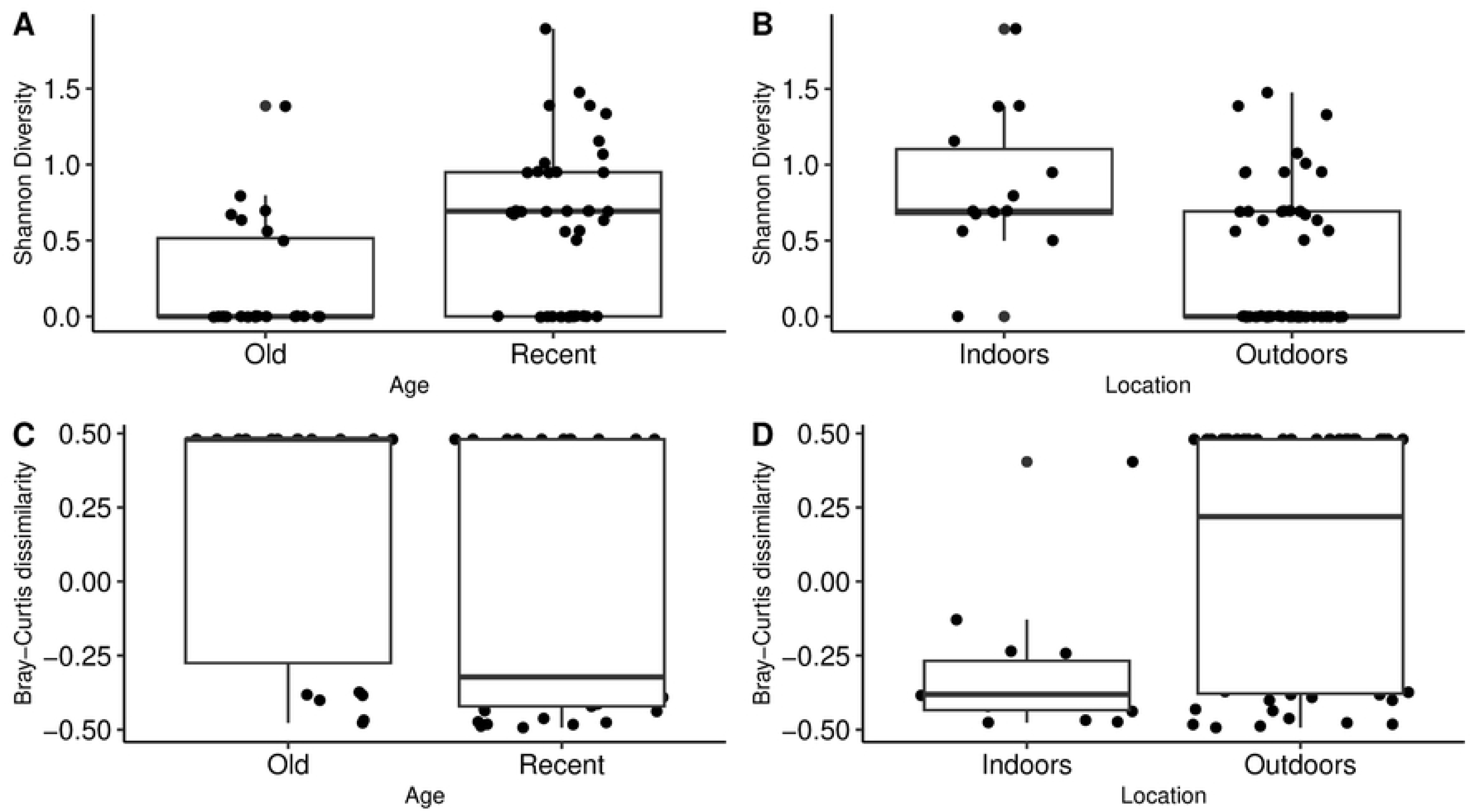
Diversity analyses of microbiome abundances. (A) Shannon diversity between old and recent samples. (B) Shannon diversity between indoor and outdoor samples. (C) Bray–Curtis dissimilarity between old and recent samples. (D) Bray–Curtis dissimilarity between indoor and outdoor samples. “Old” refers to periods prior to the post-medieval, while “recent” refers to post-medieval periods. Statistical tests: Wilcoxon rank-sum test for Shannon diversity (*p* < 0.005) and Kruskal–Wallis test for Bray–Curtis dissimilarity (*p_age_* = 0.03; *p_location_*= 0.003).

**S5 Figure.**
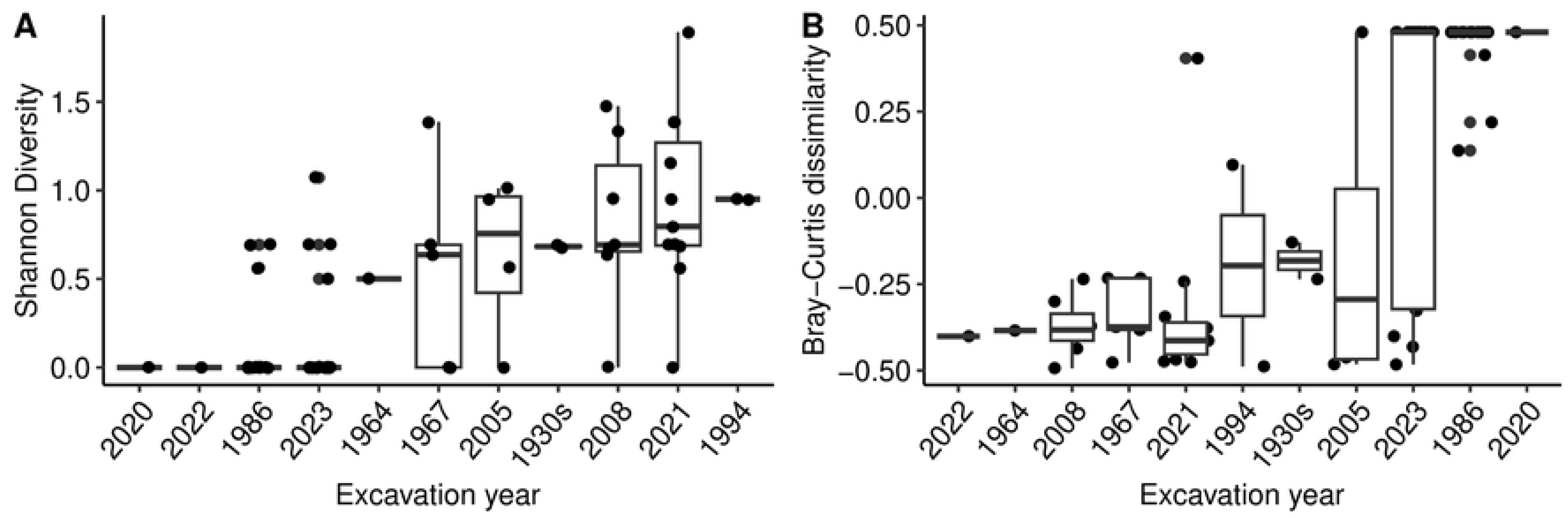
Changes in (A) Shannon diversity and (B) Bray–Curtis dissimilarity across excavation years.

**S6 Figure.**
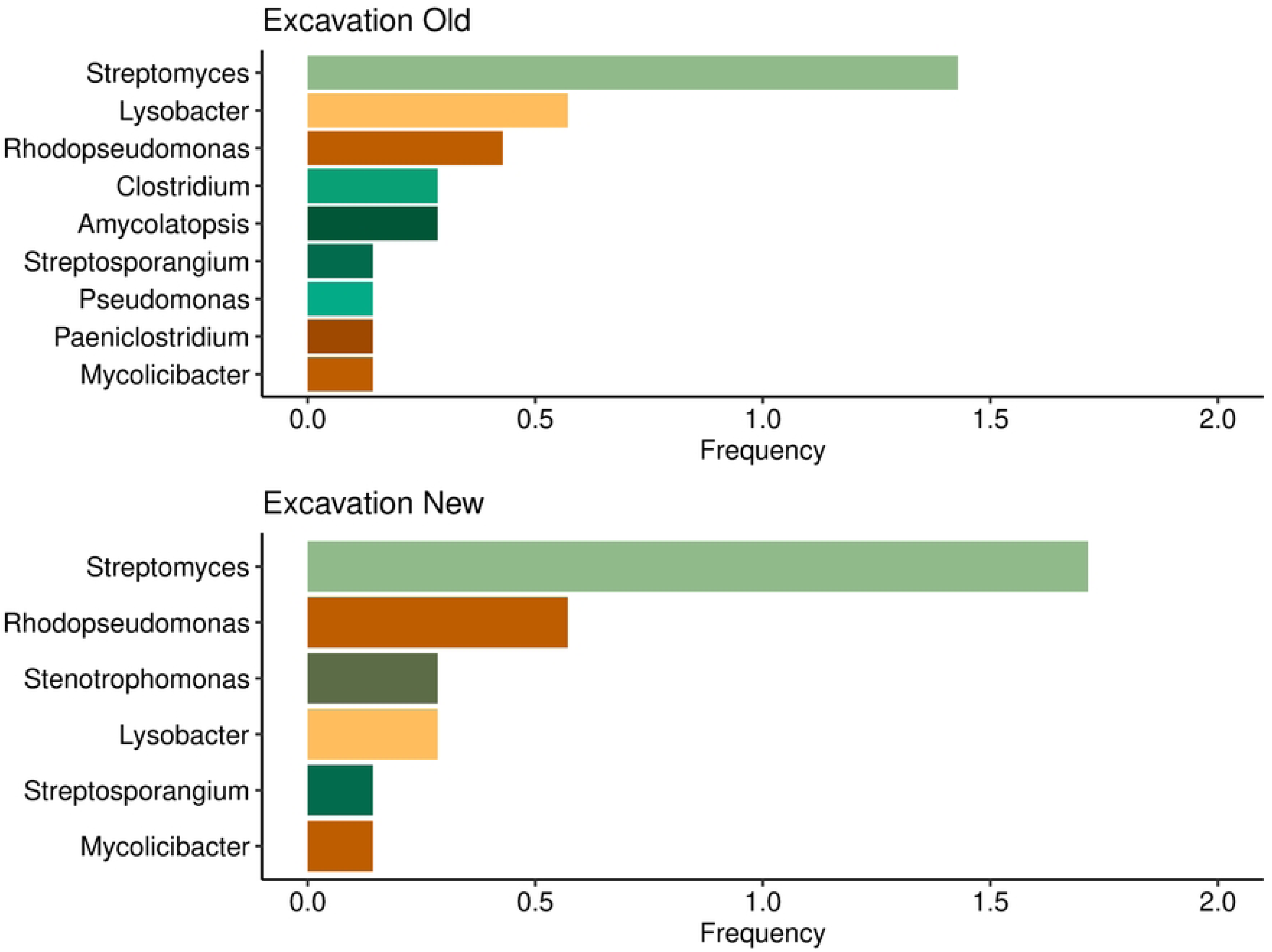
Microbiome composition of outdoor samples. Samples with only OHI scores 4 and 5, grouped into Excavation Old (1967, 1994, 2008 excavations, n=7) and Excavation New (2023 excavation, n=7).

**S7 Figure.**
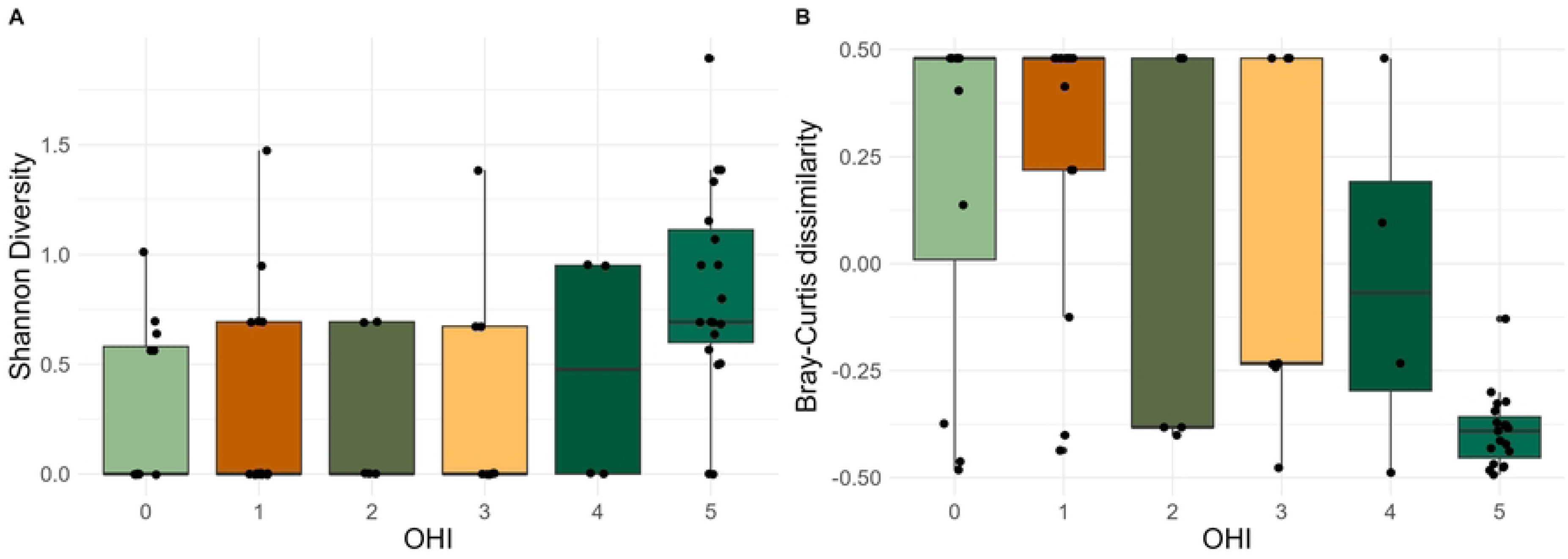
Microbial diversity analyses between different preservation scores (OHI). (A) Shannon diversity (B) Bray-Curtis diversity.

**S1 Appendix. Detailed description of excavation sites.**

**S1 Table. Basic contextual information and histological results for all samples.**

**S2 Table. KrakenUniq Abundance Matrix.**

**S3 Table. MALT Sequence statistics.**

